# Recovering Missing Regulations by Multiplex Graph Aggregation of GRN Predictions (Technical Report)

**DOI:** 10.64898/2026.11.16.767195

**Authors:** Tsz Pan Tong, Jun Pang

## Abstract

Gene regulatory networks (GRNs) capture the complex gene interactions underlying biological systems. However, existing GRN inference models face technical challenges in recovering faithful GRNs from single-cell RNA sequencing data in an unsupervised setting. GRN aggregation has been proposed to leverage the wisdom of different GRN models and improve overall performance, but aggregation methods either train on ground-truth edges, overlook the network nature of GRNs, or are computationally expensive. To bridge this gap, we propose MuxGRN, a novel GRN aggregation method that leverages multiplex link prediction to aggregate GRNs inferred from various state-of-the-art models and extract the information propagated within different layers of GRNs. We use CompGCN for representation learning and the variational graph autoencoder framework for self-supervised link prediction, which does not require ground-truth labels and can capture GRN network structure efficiently. Under this formulation, MuxGRN aggregates GRNs inferred by 9 GRN models and is evaluated on 14 real single-cell RNA sequencing datasets against the individual models and two edge-level aggregation baselines. Our evaluation shows that MuxGRN recovers up to 26.55% of previously missed true-positive edges that individual GRN models miss, substantially exceeding edge-level baselines while maintaining AUPRC comparable to the best individual GRN models. This supports the importance of the network structure in GRN and the potential for link prediction in GRN aggregation.

## 1 Introduction

Gene regulation is a core component of systems biology, characterizing the cascade of pathways that govern cell development and differentiation. Genes do not work in isolation but cooperate as a network during regulation. The modular structure, such as feedforward loops and single-input modules, serves as meaningful functional units in gene regulatory networks (GRNs), which is essential in GRN inference [20, 1]. From the perspective of network science, signaling path-ways and co-regulated genes are cascades and modules in the GRN. Hence, GRN inference involves not only predicting individual edges but also recovering network structures, which are crucial for studying underlying biological processes.

Many existing GRN inference methods often overlook the intrinsic network nature of GRNs and focus only on pairwise or triplet interactions among genes. For example, mutual information (MI) methods such as ARACNe [11], CLR [5], MINET [13], and PREMER [25] compute pairwise MI as gene-interaction statistics and use the data-processing inequality, denoising, and transfer entropy to eliminate indirect edges. Information theory methods such as PIDC [4] decompose the interaction up to three genes, while tree-based methods such as GE-NIE3 [7] and GRNBoost2 [14] only quantify the contribution of one gene in reconstructing the expression of the target gene. These methods emphasize local interactions but neglect the global network structure of GRNs. In contrast, ordinary differential equation (ODE) methods such as SCODE [12] and GRISLI [3] can capture network structures during the modeling of cell dynamics because the inferred network is multiplied iteratively during the ODE integration, and signals are propagated along the inferred network. However, their performance is not competitive with state-of-the-art (SOTA) methods [16] due to scalability issues and the assumption of linear cell dynamics. Nonetheless, the weaknesses of ODE methods should not obscure the importance of GRN network structures, and a network-aware GRN inference method remains missing.

While this may motivate the development of new network-aware GRN inference methods, we propose leveraging existing GRN inference methods developed over the years, which have proven effective in edge-level prediction, and designing a novel GRN aggregation method to combine weak signals from different GRNs. Aggregating GRN also addresses the challenge of selecting the best GRN model from a large number of bioinformatics methods, which is often difficult for biologists without computational expertise. Individual GRN models have their own strengths and weaknesses. Missing edges in one model can be captured by other models, and an aggregation method can utilize contradictory, complementary, or consistent edges, as well as the network structure of different GRNs to recover candidate regulatory edges.

Aggregating GRN is not a new idea, but existing GRN aggregation methods either require ground-truth for training [2, 19], focus on edge-level aggregation without considering the network structure [8], or optimize the aggregated graph topology using complex genetic algorithms [18]. Ground-truth labels from databases such as STRING are rare, and using them as training labels hinders independent model validation resources. Edge-level aggregation methods, such as COFFEE [8], cannot capture global network structures, whereas genetic algorithm-based methods, such as GENECI [18], are computationally expensive and converge slowly.

We recast GRN aggregation as a multiplex link prediction problem, in which GRNs from different models are treated as layers in a multiplex network, and new edges are predicted via information propagation within and across layers. GRNs at each layer can be incomplete and noisy, but the consistent edges across different layers can be reinforced, and models with complementary edges can be aggregated to discover new edges.

Motivated by this, we proposed MuxGRN, a novel GRN aggregation method that leverages CompGCN [24] for representation learning and the variational graph autoencoder (VGAE) framework for self-supervised link prediction. This empowers our method to aggregate noisy GRNs, strengthen the signals propagated across GRNs, and discover new edges that are missed in individual GRN models. Our evaluation of 14 experimental datasets on the BEELINE bench-mark [16] shows that MuxGRN can achieve performance comparable to the best individual GRN model and surpass COFFEE among the aggregation baselines. In addition, our method recovers up to 26.55% of previously missed true positive edges. This suggests that MuxGRN can exploit graph-level information for GRN aggregation. Overall, our contributions in this work are twofold:

- We propose a novel, self-supervised, and network-aware GRN aggregation method, MuxGRN, that views multiple GRNs as a multiplex network and leverages multiplex link prediction for aggregation.
- MuxGRN achieves a top-tier AUPRC among individual GRN models on 14 experimental datasets in the BEELINE benchmarks, and we empirically show that MuxGRN can recover true positive edges missed by GRN models.

These results highlight the importance of network structure in GRNs and call for future GRN inference methods to focus on their network nature.

## 2 Related Work

Marbach et al. [10] first proposed an edge-level aggregation by taking the average ranks of edges across multiple inferred GRNs, and they reported top-tier performance in the DREAM5 benchmark. COFFEE [8] introduces the use of Borda count, a rank-based voting method, for robust GRN aggregation, and the method shows a higher F-score than other individual GRN models on experimental datasets in most cases. However, rank-based methods ignore the confidence scores of each edge and cannot recover missing edges among all GRN predictions. As for topology-guided aggregation, GENECI [18] finds an optimal linear combination of GRN predictions that maximizes the topological measures of the aggregated GRN using genetic algorithms. Because graph statistics are highly sensitive to changes in edge weights, genetic algorithms are used to find optimal weights, which are computationally expensive and converge slowly. Moreover, reweighting only weakly improves performance because it does not fundamentally change relative edge strengths.

Realizing the importance of the topological structure of GRN, Fujii et al. [6] reweight the edge confidence score of GRN predictions to maximize the discrepancy between true edges and false edges using integer programming. However, identifying true and false edges leaks ground-truth information, which is not available in practice. EnsInfer [19] and EnGRaiN [2] train a meta-model using GRNs as input and part of the ground-truth edges as labels to predict the rest of the ground-truth. Although ground-truth labels are strong supervision signals, the predictions are biased towards the ground-truth and are exposed to the risk of incorrect edges in the ground-truth. Thus, a network-aware GRN aggregation method that does not rely on the ground-truth is still missing.

## 3 Preliminaries

This section formalizes the GRN aggregation task and reviews the core components, VGAE and CompGCN, used in MuxGRN.

### 3.1 Notations and Problem Definition

A GRN is a weighted directed graph *G* = (*V, ℰ, w*), where *V, ℰ* denote the sets of genes and edges, respectively, and *w* denotes the confidence score associated with each edge. In a *n*-gene system, we precompute GRN using *R* different representative GRN inference methods, each represented as a weighted adjacency matrix *A*^(*r*)^ *∈ ℝ*^*n×n*^, where *r* = 1, 2, …, *R*. Each entry 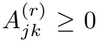 in the matrix represents the strength of regulation confidence from gene *j* to gene *k* in the GRN inferred by method *r*. For signed GRNs, we take the absolute value and discard the sign information. Our goal is to aggregate a GRN from *R* GRNs inferred by different methods without using any ground-truth labels.

### 3.2 Variational Graph Autoencoder

Variational graph autoencoder (VGAE) [9] is a self-supervised learning framework for learning node embeddings, which can be used for link prediction. Similar to a standard variational autoencoder, VGAE consists of an encoder and a decoder. The encoder *q*(*Z* | *A*) takes the adjacency matrix *A* as input and learns a latent representation *Z* of the nodes, while the decoder *p*(*A* | *Z*) reconstructs the adjacency matrix *Â* from the latent representation *Z*. To ensure a meaningful latent representation, variational inference is used such that:

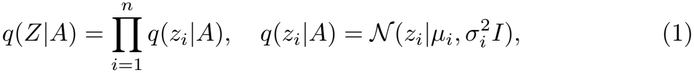

and the loss function is defined as:

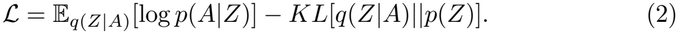

The former term is the reconstruction loss, while the latter term is the Kullback-Leibler divergence that regularizes the approximate posterior *q*(*Z*|*A*) toward the gaussian prior distribution *p*(*Z*) = Π_*i*_ *N* (*z*_*i*_|0, *I*). The adjacency matrix *Â* is reconstructed by the inner product of *Z*, i.e., *p*(*A*|*Z*) = *Â* = *σ*(*ZZ*^*T*^), where *σ* is the sigmoid function. Under this formulation, new links emerging in the reconstructed adjacency matrix *Â* can be considered potential new gene regulations.

### 3.3 CompGCN Encoder on Multiplex Networks

A multiplex network is a special type of multi-relational graph *G* = (*V, ℰ, ℛ*), where the same set of nodes *V* are connected by different layers *ℛ* of edges *ℰ*. Different GRNs inferred by different methods can be viewed as different layers in a multiplex network, and the multiplex graph allows information propagation within and across layers. In this work, we use CompGCN [24] as the encoder to learn node embeddings from the multiplex network of GRNs.

CompGCN maintains learnable node embeddings 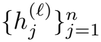 and relation embeddings 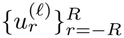 at hidden layer *ℓ*, and additionally constructs its inverse edge set *ℰ* ^*−* 1^ = *{*(*k, j, ™r*)|(*j, k, r*) *∈ ℰ}* with a new type of relation *™r*. At hidden layer *ℓ*, the message from node *j* under relation *r* is formed by element-wise multiplication 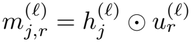.

To distinguish edge directions, messages from original and inverse edges are transformed by different linear maps 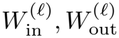 and then mean aggregated by:

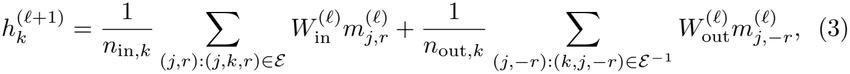

where *n*_in,*k*_, *n*_out,*k*_ count the relation-specific incoming and outgoing messages of node *k*, respectively. Relation embeddings are updated jointly by 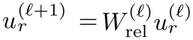. Node embedding at the final layer serves as the latent representation of the node for link prediction.

## 4 Methodology

Figure 1 describes our method, MuxGRN, which treats GRNs inferred by different representative models as distinct layers in a multiplex network, learns a shared node embedding across model layers, and reconstructs an aggregated network within the VGAE framework. New edges are predicted from node embeddings, which can be considered as potential new gene regulations.

**Fig 1.**
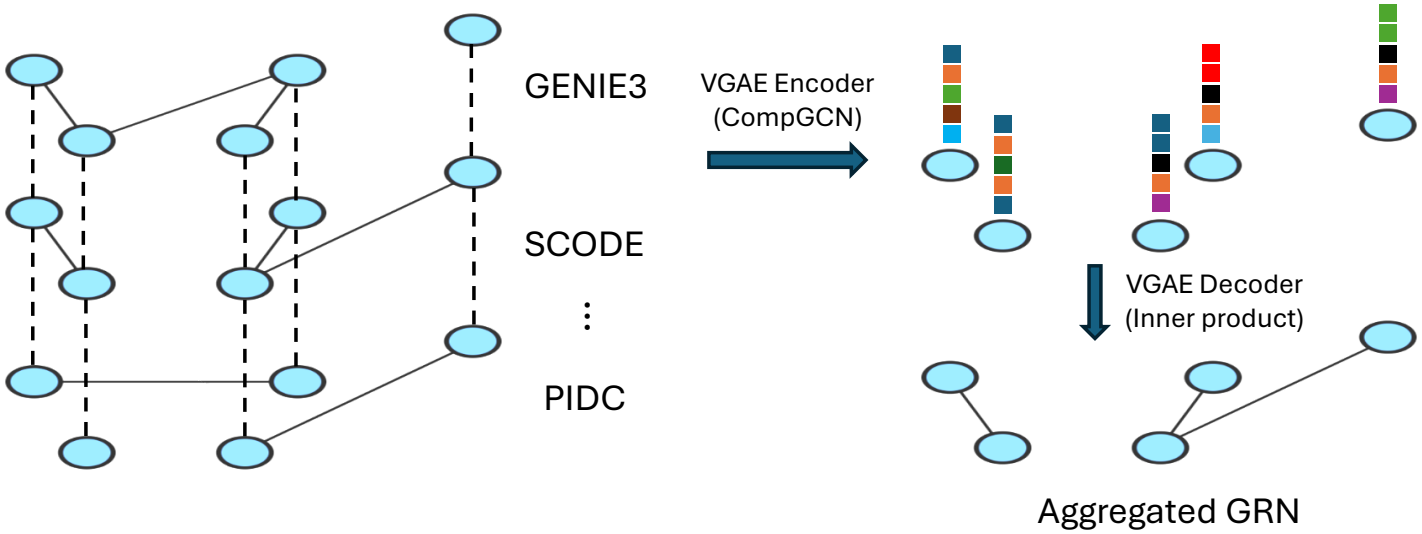
The overview of our method MuxGRN. GRNs inferred by different methods are viewed as layers in a multiplex network, and node embeddings are learned by VGAE.

We frame GRN aggregation as a multiplex link prediction problem because a multiplex graph can retain the source GRN of each edge and allow information propagation within and across different GRNs. Preserving the source GRN of each edge is essential for the aggregation because different models have different capacities to capture distinct characteristics of the underlying biological system. The edges in each pair of GRNs may be either *contradictory, complementary, or consistent*, and learning how to integrate these models via relational learning can yield a more comprehensive view of the system. In practice, given *R* GRNs inferred by different methods, we first extract the top 3*n* edges in each GRN to construct a sparse multiplex network, where 3*n* is chosen to match the empirical average degree of the ground-truth GRNs.

After that, the VGAE introduced in Section 3.2 is employed to learn the node embeddings from the multiplex network. VGAE is a self-supervised learning framework that does not require any ground-truth labels. By compressing the input multiplex network into a low-dimensional latent space and reconstructing an aggregated network from the latent representation, VGAE can distill the essential information from the noisy GRNs and capture intrinsic network structures in the multiplex network. Unreliable edges in the input GRNs can be filtered out during compression, consistent edges across different GRNs can be reinforced, and missing edges will emerge in the reconstructed multiplex network.

Motivated by this intuition, CompGCN introduced in Section 3.3 is chosen as the encoder to learn node embeddings and relation embeddings from the multiplex network. CompGCN is specially designed for multi-relational graphs, which can learn node and relation representations simultaneously. Since the relation represents each GRN model, the relation embedding captures the characteristics of each GRN model, decoupling the model-specific information from the node embedding and allowing the node embedding to capture only the shared information across different GRNs.

Finally, this consensus information is then used to reconstruct the network using the inner product decoder. Because of the information propagation on the multiplex network, the reconstructed edges are no longer an edge-level aggregation of the input GRNs but carry graph contextual information. In edge aggregation methods, only consistent edges across different GRNs are reinforced, contradictory edges are diminished, and complementary edges are weakened. In contrast, our method can capture both consistent and complementary signals through relation learning, which helps recover missing edges. Detailed implementation is provided in our public code repository https://github.com/1250326/MuxGRN.

## 5 Experiments

This section reports the evaluation pipeline and assesses whether MuxGRN can recover new edges that its source models miss while preserving overall predictive performance.

### 5.1 Datasets and evaluation metrics

We evaluated MuxGRN on the BEELINE benchmark [16], which contains 7 sets of experimental scRNA-seq datasets for human and mouse biological systems, namely, hESC, hHep, mDC, mESC, mHSC-E, mHSC-GM, and mHSC-L. The top 500 and top 1,000 high-variance genes and their transcription factors are selected from each dataset, resulting in 14 datasets in total with 501–1,783 genes. All available functional-association edges in the STRING database are used to construct the undirected ground-truth GRNs. Depending on the gene sets in different datasets, ground-truth GRNs have 3,220–85,424 edges and 0.92%–4.19% density. Note that the ground-truth GRNs are not used for training.

We evaluated the performance of all benchmarking models using AUPRC to measure their precision and discriminative ability. We also computed the proportion of newly discovered true positive edges apart from edges in the union of all individual GRN models, normalized by the best individual GRN models for aggregation baselines to show the ability to find new edges. Denote *E*_Agg_, *E*_*r*_, *E*_true_ as the sets of predicted edges in an aggregation model Agg, an individual model *r*, and the ground-truth, respectively. The proportion is calculated as:

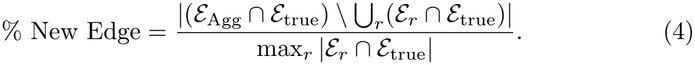

All reported results are averaged over 10 random initializations.

### 5.2 Representative GRN models and aggregation baselines

Nine representative GRN inference methods from BEELINE and its subsequent GRN benchmarks are selected from different categories, including correlation (LEAP [21]), information theory (PIDC [4], Scribe [17]), tree methods (GE-NIE3 [7], GRNBoost2 [14], FlowGRN [23]), regression (SINCERITIES [15]), and ODE methods (SCODE [12], GRISLI [3]). Methods requiring prior GRN knowledge or exceeding the computational budget on these datasets were excluded. GRNs are inferred using the default parameters in BEELINE, then aggregated by MuxGRN and two edge-level aggregation baselines, COFFEE [8] and Mean.

COFFEE uses the Borda count to compute the edge rank in each GRN and averages the rank score across different GRNs. Denote the rank of edge (*j, k*) in GRN *A*^(*r*)^ as rank 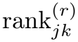, the aggregated GRN is calculated as:

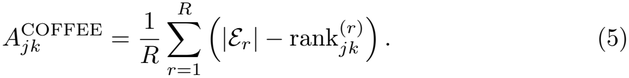

In addition, we designed a mean aggregation to compute the z-score of the edge weights for each GRN, and then averaged the z-scores across GRNs. Denote the mean and standard deviation of edge weights in GRN *A*^(*r*)^ as *µ*^(*r*)^ and *σ*^(*r*)^, respectively, the aggregated GRN is calculated as:

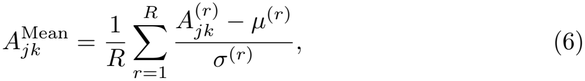

### 5.3 Implementation details

For MuxGRN, our VGAE used a two-layer CompGCN as the encoder and inner product as the decoder. We set the input, hidden, and output embedding dimensions to 1, 64, and 64, respectively, for the encoder, with multiplication as the composition operation and ReLU as the activation function. The model is trained for 1,000 epochs using the Adam optimizer with a learning rate of 0.01 and a negative sampling ratio of 1.0. Standard binary cross entropy and the Kullback-Leibler divergence are used as the loss functions. Our model is trained on a single NVIDIA V100 SXM2 GPU with 16 GB of memory, and the training time is below 1 minute for all datasets. Since STRING ground-truths are undirected, all directed predictions are symmetrized before evaluation.

### 5.4 Evaluation results

Figure 2 shows the average AUPRC and the proportion of newly discovered true-positive edges of different GRN models on 14 BEELINE datasets across 10 random initializations. MuxGRN achieves a similar AUPRC to the top-tier individual GRN models. Compared with two edge-level aggregation baselines, MuxGRN achieves a similar AUPRC to the mean baseline and outperforms COFFEE on 13 of 14 datasets. Our method also discovers up to 26.55% of previously missed true-positive edges, surpassing the maximum of 9.50% for COFFEE and 0.33% for Mean. This shows that MuxGRN can discover true edges that individual GRN models miss while maintaining a top-tier AUPRC.

**Fig 2.**
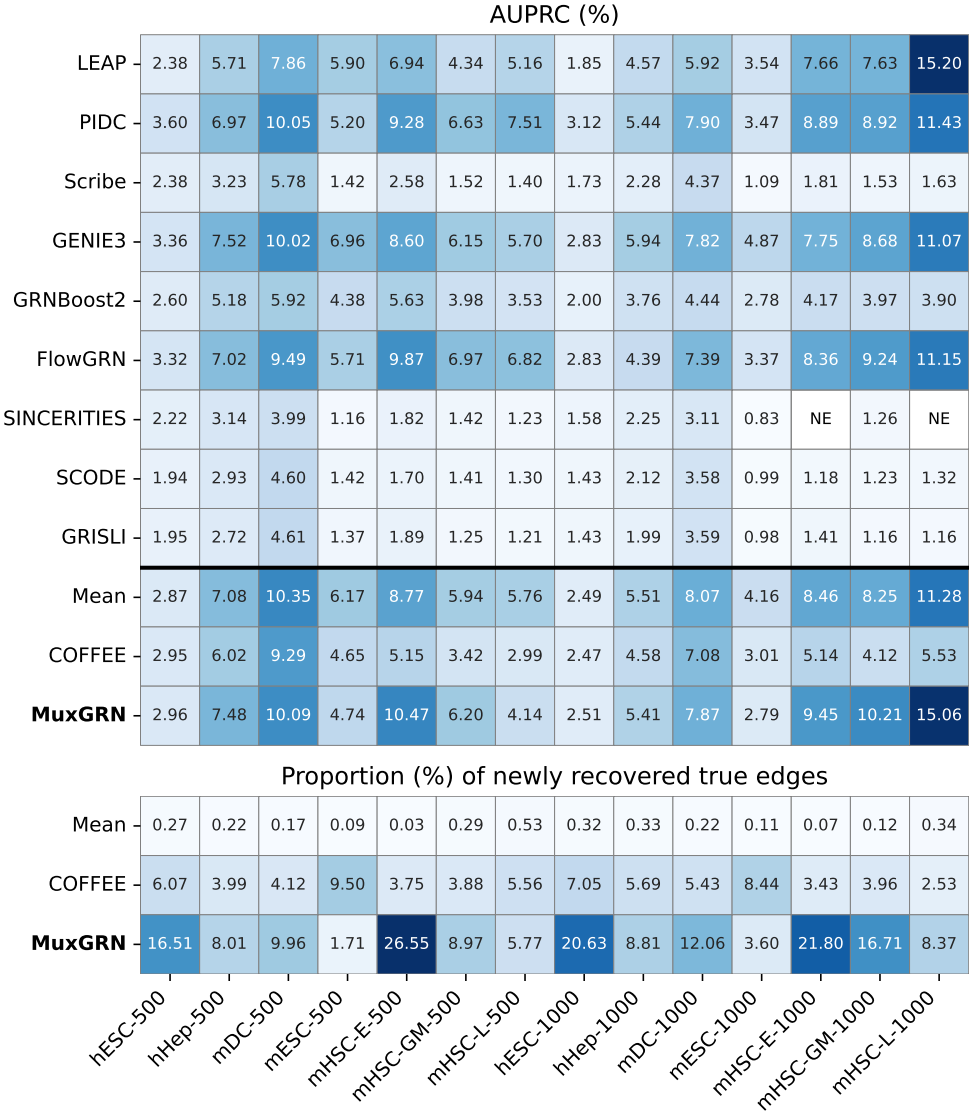
Average AUPRC (%) (upper) and average proportion (%) of newly discovered true-positive edges (lower) of different GRN models on 14 BEELINE datasets over 10 random initializations. “NE” indicates numerical error.

## 6 Conclusion and Limitations

Different GRN inference methods have different strengths and weaknesses in capturing the underlying biological system, and thus their inferred GRNs may be contradictory, complementary, or consistent. Manually integrating these GRNs or choosing one of them is infeasible and suboptimal in large networks with thousands of genes, thus a systematic GRN aggregation method is required to resolve different GRN predictions and recover candidate regulatory edges.

In this work, we designed a novel network-aware GRN aggregation method, MuxGRN, to extract weak signals from different heterogeneous GRNs through multiplex link prediction. MuxGRN is designed for settings where no ground-truth GRN is available to select the retrospectively best individual method. This vision is also supported by our evaluation, which shows that graph-level aggregation can recover previously missed true edges while preserving competitive global edge precision. Results indicate that network-level aggregation can recover additional interactions without relying on ground-truth labels during training. Our findings also call for more network-aware GRN inference methods that can capture the modular structure of GRNs, which is essential for understanding the underlying biological system.

Despite promising results, our method has several limitations. First, Mux-GRN currently does not infer regulatory direction or sign because the STRING-based reference is undirected and the inner-product decoder is symmetric. Besides, the sensitivity of MuxGRN to the selected source-model panel remains an open question. Furthermore, MuxGRN lacks model explainability because it does not learn explicit scalar weights for source GRNs, and source-specific information is encoded implicitly through relation embeddings in CompGCN. Future work focuses on overcoming these limitations.

## Acknowledgments

Authors Tsz Pan Tong and Jun Pang acknowledge financial supports from the Luxembourg National Research Fund (FNR) under the grant agreement INTER/NCN/24/18732364/EdgeCR, and the Institute for Advanced Studies of the University of Luxembourg through an Audacity Grant (AUDACITY-2021). A preliminary two-page version of this work was presented orally and published as an extended abstract at GIW XXXIV ISCB-Asia 2025 [22].

## A Appendix

Figure 3 reports the standard deviation of AUPRC (%) and the proportion (%) of newly discovered true-positive edges in the benchmark. MuxGRN has the highest deviations in both metrics due to the stochasticity of the deep learning training process. However, the magnitude of the deviations does not alter the conclusion on the advantages of MUXGRN over other aggregation baselines.

**Fig 3.**
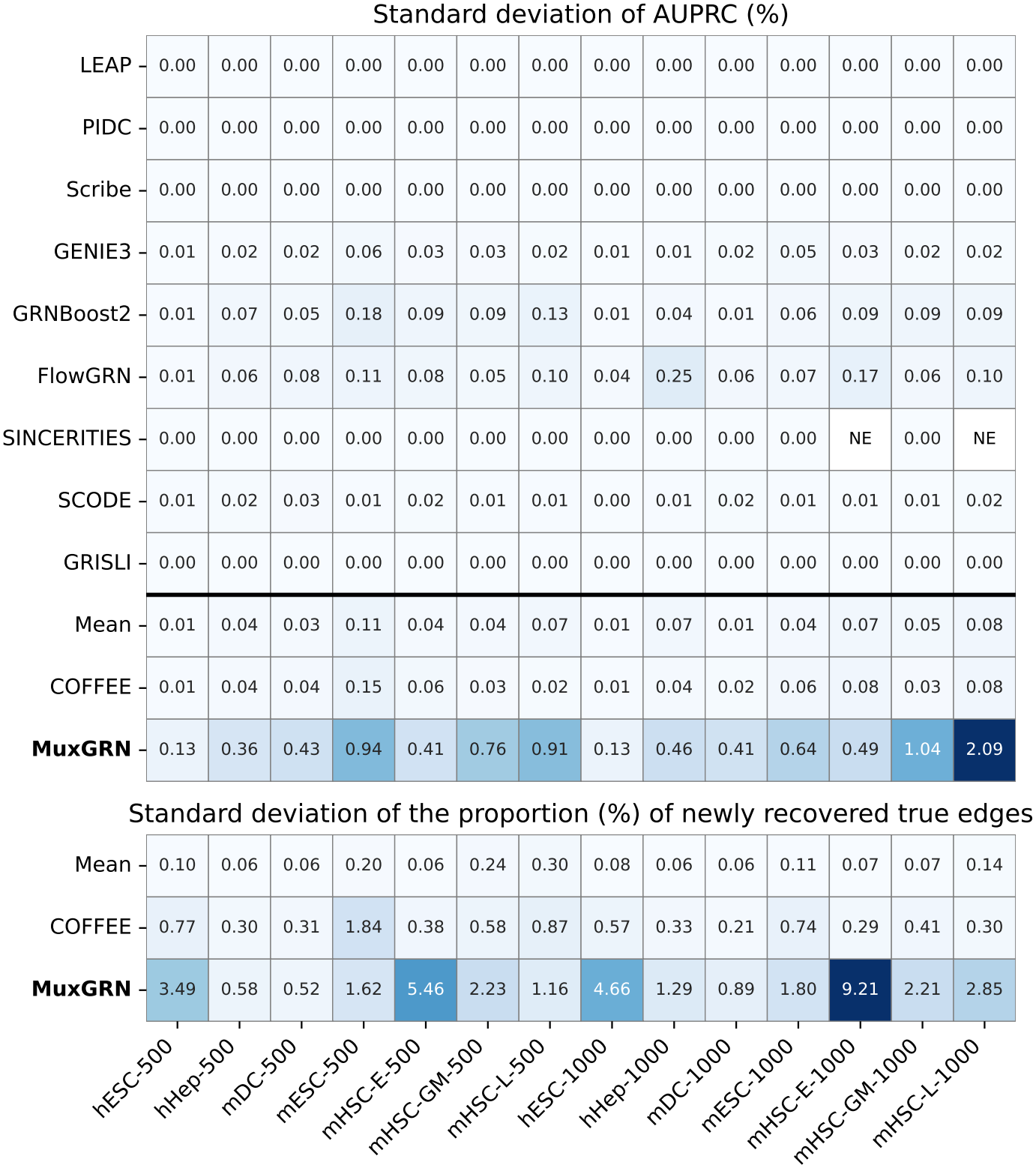
Standard deviation of AUPRC (%) (upper) and proportion (%) of newly discovered true-positive edges (lower) of models on the BEELINE benchmark.

## References

1. Alon, U.: Network motifs: theory and experimental approaches. Nature Reviews Genetics 8(6), 450–461 (2007). 10.1038/nrg2102

2. Aluru, M., Shrivastava, H., Chockalingam, S.P., Shivakumar, S., Aluru, S.: EnGRaiN: a supervised ensemble learning method for recovery of largescale gene regulatory networks. Bioinformatics 38(5), 1312–1319 (2022). 10.1093/bioinformatics/btab829

3. Aubin-Frankowski, P.C., Vert, J.P.: Gene regulation inference from single-cell rna-seq data with linear differential equations and velocity inference. Bioinformatics 36(18), 4774–4780 (2020). 10.1093/bioinformatics/btaa576

4. Chan, T.E., Stumpf, M.P., Babtie, A.C.: Gene Regulatory Network Inference from Single-Cell Data Using Multivariate Information Measures. Cell Systems 5(3), 251–267 (2017). 10.1016/j.cels.2017.08.014

5. Faith, J.J., Hayete, B., Thaden, J.T., Mogno, I., Wierzbowski, J., Cottarel, G., Kasif, S., Collins, J.J., Gardner, T.S.: Large-Scale Mapping and Validation of Escherichia coli Transcriptional Regulation from a Compendium of Expression Profiles. PLOS Biology 5(1), e8 (2007). 10.1371/journal.pbio.0050008

6. Fujii, C., Kuwahara, H., Yu, G., Guo, L., Gao, X.: Learning gene regulatory networks from gene expression data using weighted consensus. Neurocomputing 220, 23–33 (2017). 10.1016/j.neucom.2016.02.087

7. Huynh-Thu, V.A., Irrthum, A., Wehenkel, L., Geurts, P.: Inferring regulatory networks from expression data using tree-based methods. PLOS ONE 5(9), e12776 (2010). 10.1371/journal.pone.0012776

8. K Lodi M., Chernikov, A., Ghosh, P.: COFFEE: consensus single cell-type specific inference for gene regulatory networks. Briefings in Bioinformatics 25(6), bbae457 (2024). 10.1093/bib/bbae457

9. Kipf, T.N., Welling, M.: Variational graph auto-encoders. arXiv preprint arXiv:1611.07308 (2016). 10.48550/arXiv.1611.07308

10. Marbach, D., Costello, J.C., Küffner, R., Vega, N.M., Prill, R.J., Camacho, D.M., Allison, K.R., Kellis, M., Collins, J.J., et al.: Wisdom of crowds for robust gene network inference. Nature Methods 9(8), 796–804 (2012). 10.1038/nmeth.2016

11. Margolin, A.A., Nemenman, I., Basso, K., Wiggins, C., Stolovitzky, G., Favera, R.D., Califano, A.: ARACNE: An algorithm for the reconstruction of gene regulatory networks in a mammalian cellular context. BMC Bioinformatics 7(1), S7 (2006). 10.1186/1471-2105-7-S1-S7

12. Matsumoto, H., Kiryu, H., Furusawa, C., Ko, M.S., Ko, S.B., Gouda, N., Hayashi, T., Nikaido, I.: SCODE: an efficient regulatory network inference algorithm from single-cell RNA-Seq during differentiation. Bioinformatics 33(15), 2314–2321 (2017). 10.1093/bioinformatics/btx194

13. Meyer, P.E., Lafitte, F., Bontempi, G.: MINET: A R/Bioconductor package for inferring large transcriptional networks using mutual information. BMC Bioinformatics 9(1), 461 (2008). 10.1186/1471-2105-9-461

14. Moerman, T., Aibar Santos, S., Bravo González-Blas, C., Simm, J., Moreau, Y., Aerts, J., Aerts, S.: GRNBoost2 and Arboreto: efficient and scalable inference of gene regulatory networks. Bioinformatics 35(12), 2159–2161 (2019). 10.1093/bioinformatics/bty916

15. Papili Gao, N., Ud-Dean, S.M., Gandrillon, O., Gunawan, R.: SINCERITIES: inferring gene regulatory networks from time-stamped single cell transcriptional expression profiles. Bioinformatics 34(2), 258–266 (2018). 10.1093/bioinformatics/btx575

16. Pratapa, A., Jalihal, A.P., Law, J.N., Bharadwaj, A., Murali, T.: Benchmarking algorithms for gene regulatory network inference from single-cell transcriptomic data. Nature Methods 17(2), 147–154 (2020). 10.1038/s41592-019-0690-6

17. Qiu, X., Rahimzamani, A., Wang, L., Ren, B., Mao, Q., Durham, T., McFaline-Figueroa, J.L., Saunders, L., Trapnell, C., Kannan, S.: Inferring causal gene regulatory networks from coupled single-cell expression dynamics using Scribe. Cell Systems 10(3), 265–274 (2020). 10.1016/j.cels.2020.02.003

18. Segura-Ortiz, A., García-Nieto, J., Aldana-Montes, J.F., Navas-Delgado, I.: GENECI: a novel evolutionary machine learning consensus-based approach for the inference of gene regulatory networks. Computers in Biology and Medicine 155, 106653 (2023). 10.1016/j.compbiomed.2023.106653

19. Shen, B., Coruzzi, G., Shasha, D.: EnsInfer: a simple ensemble approach to network inference outperforms any single method. BMC Bioinformatics 24(1), 114 (2023). 10.1186/s12859-023-05231-1

20. Shen-Orr, S.S., Milo, R., Mangan, S., Alon, U.: Network motifs in the transcriptional regulation network of escherichia coli. Nature genetics 31(1), 64–68 (2002). 10.1038/ng881

21. Specht, A.T., Li, J.: LEAP: constructing gene co-expression networks for single-cell RNA-sequencing data using pseudotime ordering. Bioinformatics 33(5), 764–766 (2017). 10.1093/bioinformatics/btw729

22. Tong, T.P., Pang, J.: Aggregating GRNs via multiplex link prediction. Briefings in Bioinformatics 26(Supplement_1), i11–i12 (2025). 10.1093/bib/bbaf631.012

23. Tong, T.P., Pang, J.: FlowGRN: Scalable and dropout-robust gene regulatory network inference via flow matching-based trajectory reconstruction. In: Proceedings of the 16th ACM International Conference on Bioinformatics, Computational Biology, and Health Informatics. pp. 1–8 (2025). 10.1145/3765612.3767196

24. Vashishth, S., Sanyal, S., Nitin, V., Talukdar, P.: Composition-based multi-relational graph convolutional networks. In: Proceedings of the International Conference on Learning Representations (2020). 10.48550/arXiv.1911.03082

25. Villaverde, A.F., Becker, K., Banga, J.R.: PREMER: a tool to infer biological networks. IEEE/ACM Transactions on Computational Biology and Bioinformatics 15(4), 1193–1202 (2017). 10.1109/TCBB.2017.2758786

